# Infectious Bronchitis: Molecular Detection and Characterization in Live Bird Markets in Jos, Plateau State, Nigeria

**DOI:** 10.1101/2025.03.02.641078

**Authors:** Bitrus Inuwa, Hassan Ismail Musa, Hambali Ibrahim Umar, Abdulyeken Olawale Tijjani, Shuaibu Gidado Adamu, Muhammad Abubakar Sadiq, Ismaila Shittu, Clement Meseko, Tony Joannis

## Abstract

Infectious bronchitis virus (IBV), the avian coronavirus, is a highly contagious coronavirus of birds. It mostly affects the respiratory, urinary and reproductive tract, leading to considerable economic losses to the poultry industry as a result of drop in egg production, undesirable egg quality and poor weight gain. Thus, this study aimed to detect and characterize the virus in live bird markets (LBMs) in Plateau State, Nigeria. One hundred pooled of cloacal and tracheal swab samples each were collected from two LBMs in Jos. Viral RNA was extracted and screened for IBV using real-time RT-PCR. Subsequently, hyper-variable region of the spike (*S1*) gene of positive samples were amplified and sequenced. IBV nucleic acid was detected in 42% (42/100) of the pooled samples. Phylogenetic analysis of the resulting sequences of *S1* gene showed that IBV detected in this study were diversified into two distinct genotypes: GI-14, and GI-23. These genotypes were closely related to a Poland strain with over 95% nucleotide identity, forming a common cluster within the GI-23 group. One isolate showed a notable relationship with a previously reported Nigerian isolate, sharing 85% nucleotide identity, and formed a common cluster within the GI-14 group. Overall, this study established the widespread of IBV and therefore recommends continuous surveillance to identify the current circulating strain for possible local vaccine development for effective control measures to mitigate the spread of the virus in the study area and the country at large.

## Introduction

The world population is growing and the demand for good-quality livestock products is expanding and expected to keep increasing. In Nigeria, over 13 million households keep livestock and receive at least part of their income from it (FAO, 2018). Nigeria currently has the second largest poultry population in Africa, with a standing stock of about 180 million birds. Annually, 454.000 tonnes of meat and more than 14 billion eggs are produced (FAO, 2018). The poultry sector contributes 6-8% of gross domestic product, which is about 30% of the total agriculture contribution (FAO, 2018). However, infectious diseases, including the coronavirus, represent a constant challenge for the poultry industry globally, causing enormous economic losses for farmers due to decreased production. Coronaviruses (CoVs) are enveloped, positive-sense, single-stranded RNA viruses belonging to family *Coronaviridae*. The coronaviruses are comprised of four genera, this include: *Alpha, Beta, Gamma*, and *Deltacoronavirus* (Cui et al., 2019; Mostafa *et al*., 2018; Inuwa *et al*., 2021). Avian coronaviruses belong to *Gammacoronavirus* genus, *Igacovirus* subgenus, subfamily *Coronavirinae*, and contain infectious bronchitis virus (IBV) and other bird coronaviruses. Avian coronavirus can cause diseases in Galliformes birds, such as chickens, turkeys, peafowl, pheasants, partridge, quails, guinea fowl, and wild birds with respiratory signs, as well as kidney diseases, undesirable egg quality, and mortality in some cases (Jackwood and De Wit, 2019). The destruction of the mucosa after infection allows the invasion of secondary pathogens, exacerbating the disease (Rautenschein & Philipp, 2021). The most prominent representative of *Gammacoronavirus* genus is IBV (Cavanagh *et al*., 2002). It is the longest known coronavirus in the world as it was first discovered by Hawk and Schawn over 90 years ago (Rautenschein and Philipp, 2021; Schalk and Hawn, 1931; Finger *et al*, 2025). IBV is detected in almost all regions of the world (Bande *et al*., 2017). Some variants appear regionally restricted and only for a short time, while others persist and spread (Legnardi *et al*., 2020). IBV is a highly contagious viral pathogen of chickens and incurs economic losses to the poultry industry globally (Cavanagh *et al*., 2002). IBV is ranked by World Organization for Animal Health as a disease causing the second largest loss in poultry production after highly pathogenic avian influenza (HPAI) followed by low pathogenic avian influenza (LPAI), Newcastle disease (NCD), and Infectious bursal disease (IBD) (Rana *et al*., 2021; Sjaak de *et al*., 2011 World Livest. Dis. Atlas, 2011). The viral genome comprises a single-stranded positive-sense RNA having a size approximately 27.5 to 28 Kb, which encodes four structural proteins that include: spike (S), envelope (E), membrane (M), nucleocapsid (N), and several non-structural proteins (Jackwood and De Wit, 2019). The spike protein is of specific interest for the virus-host interaction, as it forms projections on the virions. The spike (S) protein comprises of two subunits: S1 and S2. The S1 forms the extracellular part of the virus and plays a key role in tissue tropism, induction of protective immunity, virus neutralization, cell attachment and serotype specificity, whereas the S2 subunit anchors the spike into the virus membrane. During IBV replication and evolution, the high mutation rate within the S1 gene generates substantial genotypical, antigenic and pathogenic variations. Many newly emerging serotypes referred to as ‘‘variants’’ have been identified due to point mutations, genetic recombination events, and selective pressure in the hyper-variable regions of the genome (Jackwood and De Wit, 2019; Legnardi *et al*., 2020). Several circulating serotypes of the virus which may not confer cross-protection on each other have been identified in poultry worldwide. The unified IBV classification, based on the diversity of the S1 coding region, introduced in 2016, includes at least 32 different viral lineages arranged into six genotypes and a number of unique variants, but the identification of new genotypes/lineages was subsequently reported (Domanska-blicharz *et al*., 2014; Valastro *et al*., 2016). The continuous evolution of IBV variants in various regions across the globe remains a serious concern for poultry production globally (Domanska-blicharz *et al*., 2014; Fraga *et al*., 2018). In Nigeria, Adene and Ojo (1976) described and documented the first evidence of suspected IB disease. Previous data on IBV in Nigeria focused on seroprevalence and molecular detection (Bitrus *et al*., 2020; Shittu *et al*., 2019). There is limited information on the molecular characterization of the circulating genotype of the virus in Nigeria. This study aimed at detection and characterization of IBV from live bird markets (LBMs) in Jos, Plateau State, Nigeria. LBMs are the popular places all over the world that receive live poultry to be resold or slaughtered on-site. These markets have been identified as providing enabling environment for the dissemination of infectious diseases among bird species and potential reassortment because of the high population densities and the mixture of poultry species, ornamental and wild birds that usually co-exist. These markets are usually located near residential areas in the cities (Senne *et al*., 2003). As illustrated by the results of some epidemiological studies, not only do the LBMs play a key role in spreading infectious diseases among bird species, but they can also transfer some diseases, such as Avian Influenza, to humans (Zhou *et al*., 2015).

## Materials and Methods

### Study Area

The study was carried out in Jos the capital of Plateau State located between Lat. 9° 56’N and 8° 53’E. Jos has a near temperate climate with an average temperature between 18 and 22°C which favors poultry production, therefore making it a major producer of poultry products in the North-Central region of Nigeria.

### Sample collection

In the present study, cross-sectional survey was carried out from September 2019 to January 2021 in two live-bird markets (LBMs), Jos Plateau State, Nigeria. This two LBMs, Kugiya, located in the Southern part while Yankaji located in Northern part of Jos, were selected because there the two major central LBM that received poultry within and outside the state. A total of 100 pooled tracheal and cloacal swabs each were randomly collected from different species of birds (broiler *n* = 20, duck *n* = 50 and chicken *n* = 30) in a viral transport medium from the two LBMs. Samples were transported in a cool chain to the National Veterinary Research Institute, Vom, Nigeria, and were kept at -80 ° C until analysis.

### RNA Extraction and Detection of IBV by Real-time RT-PCR

Viral RNA was extracted directly from 140 μl of each sample pools using a QIAamp Viral RNA Mini Kit (Qiagen, Valencia, CA, USA) according to the manufacturer’s instructions. Extracted RNA was stored at -80 °C until further analysis. The RNAs were tested using QuantiTect Multiplex RT-PCR Kit (Qiagen, Germany) as previously reported (Callison *et al*., 2006; Bitrus *et al*., 2020). Primers and Taqman® hydrolysis probe targeting the conserved regions of 5’ UTR gene of the avian coronavirus genome was carried out under the following thermal cycling condition: 50°C for 30 min for RT reaction, initial denaturation at 95°C for 15 min, followed by 40 cycles at 94°C for 10 sec denaturation and 60°C at 60 sec annealing and data collection. The real-time RT-PCR amplification was performed using Rotor Gene Q (Qiagen Co., Hilden, Germany) real-time PCR system.

### Detection of IBV by partial amplification of the spike (S1) gene by conventional RT-PCR

A region corresponding to approximately 300 base pairs of partial hyper-variable region of spike gene was amplified by conventional RT-PCR using specific primer as described previously (Valastro *et al*., 2016). The PCR reaction was carried out using One-Step RT-PCR Kit (Qiagen, Hilden, Germany) in a 25 μl reaction mixture containing 5.0 μl of 5× PCR buffer, 1.0 μl dNTP mix (10 mM each), 0.5 μl of each primer (50 μM), 0.2 μl RNase inhibitor (40 U/μl, Promega), 1.0 μl RT-PCR enzyme mix, 5.0 μl of RNA template and 14.3 μl nuclease-free water to make up to the final volume. The following cycling profile were used: 50°C at 30 min, 94°C for 15 min; 40 cycles of 94°C for 30 sec, 45°C for 1 min, 68°C for 1 min, and a final extension at 68°C for 7 min. Amplification was carried out in Touch Thermal Cycler (BIO-RAD C1000). All Amplified PCR products alongside 100 base pair DNA molecular weight marker were analysed by gel electrophoresis on 1.5% agarose gel stained with ethidium bromide and visualized in Biostep Dark Hood DH-40/50 imaging analysis system (Biostec-Fischer, Germany).

### Sequencing of Partial S1 Gene and Bioinformatics Analysis

To establish the genetic relatedness of the positive samples, the amplified PCR products of all positive samples were purified using Wizard® SV Geland PCR Clean-up System (Promega) following the manufacturer’s instructions and sent for sequencing at a commercial sequencing facility (Macrogen Europe, The Netherlands). The qualities of the obtained sequences were analyzed using BioEdit (version 7.2.5.0). Using the search engine at the National Centre for Biotechnology Information (NCBI, USA) site (http://www.ncbi.nih.gov), BLAST of the sequences was carried out to obtain related IBV sequences for comparison. Alignment was performed using MUSCLE MEGA X. Phylogenetic tree of sequences with other reference IBV strains obtained from the GenBank database was constructed using the neighbor-joining method in MEGA X (Kumar *et al*., 2018). A bootstrap resampling analysis was performed (1000 replicates) to test the robustness of the major phylogenetic groups.

## Data Analysis

Descriptive statistics such as frequency and percentages were employed to present an estimate of the prevalence. Statistical analysis was performed using the statistical package for social sciences (SPSS). All P-values less than 0.05 were considered to be statistically significant.

## Result

In all, a prevalence of 42% (42/100) was recorded. At the market level, the prevalence of 37.5% (15/40) for Kugiya LBM and 45% (27/60) for Yankaji LBM were reported respectively (Table I). There was no significant association between IBV and LBMs. Positive samples were recorded in all species of birds (broiler, chicken and duck) sampled. Higher prevalence of 44% (22/50) was recorded in the duck. Both broiler and local chicken had 40% (8/20) and 40% (12/30) prevalence (Table II). More positive 25% (27/100) were recorded from the cloacal samples than the tracheal 15% (15/100). No significant association (P > 0.05) was observed between IBV and the route of sampling (Table II). There was no significant difference between IBV and the age of birds. More positive samples (61.8%) were recorded in the adult birds (above 12 weeks) than in the young birds (53.1%) in both LBMs (Table II).

**Table I:** Sample Distribution and IBV Prevalence (%) in two LBMs in Jos using Real-time RT-PCR.

| LBM Market | No. Sampled | No. Positive | % |
| --- | --- | --- | --- |
| Kugiya | 40 | 15 | 37.5 |
| Yankaji | 60 | 27 | 45.0 |
| **Total** | **100** | **42** |  |

**Table II:**
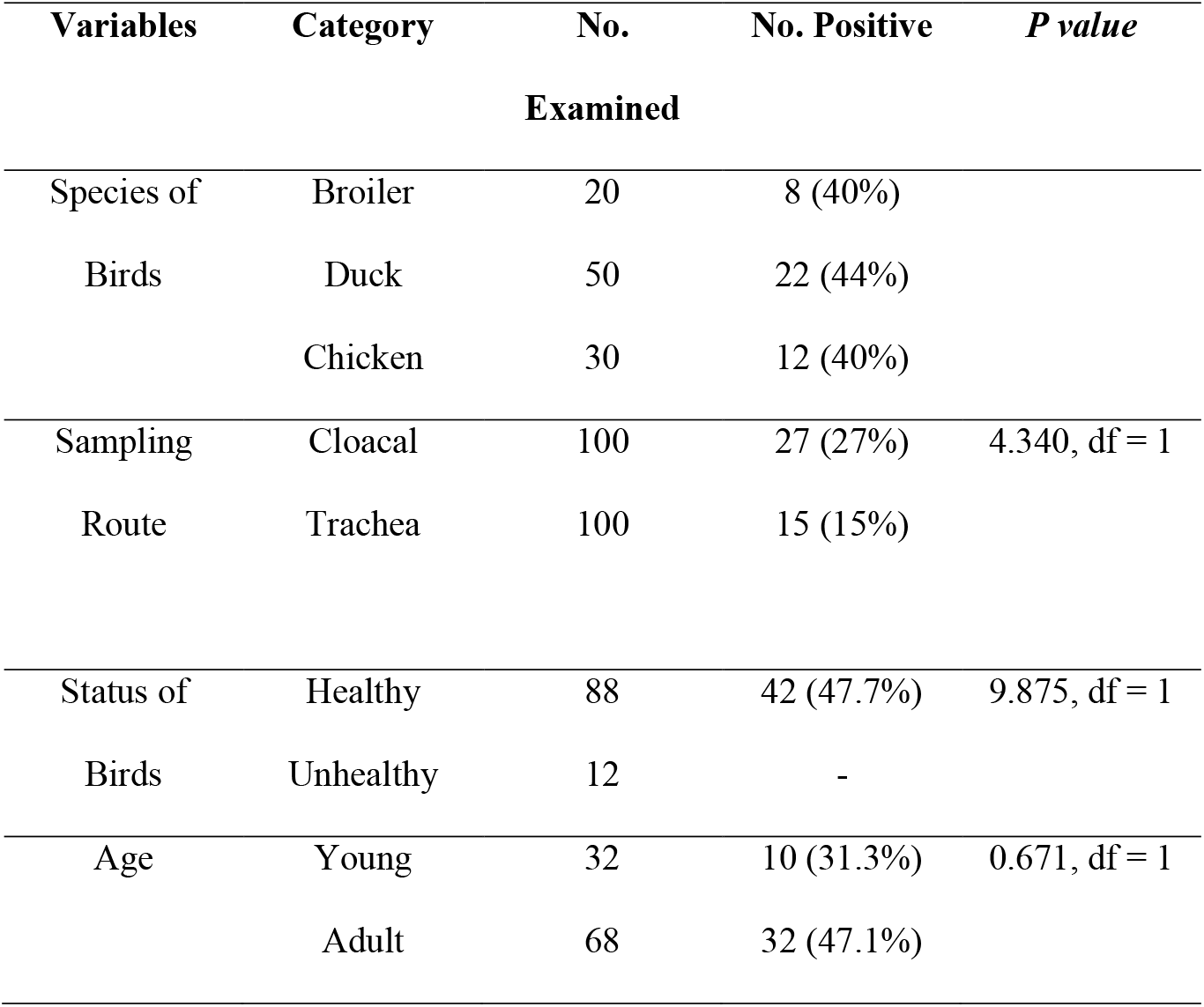
Descriptive Statistics of Sampled Birds.

### Result of Molecular Detection of the Virus using Conventional RT-PCR (*S1 gene*)

Infectious bronchitis virus was detected by the *S1* gene in four (63.6%) out of seven samples examined.. The result of the gel electrophoresis of the amplified products showed approximately 300 base pair (bp) using the *S1* glycoprotein gene primer. One hundred base pair (bp) gene marker (ladder), showed appearance of bands which correspond to the targeted position of the DNA gene marker, indicating positive IBV amplicon. All negative controls were negative using conventional RT-PCR.

### Phylogenetic Analysis Result

The result of the phylogenetic analysis of the partial nucleotide sequencing of the *S1* gene in this study indicated that the Nigerian IBV virus were diversified into two distinct genotypes: GI-14, and GI-23 (Figures I) according to (Valastro *et al*., 2016): IBV/PL/CK/17-20T/2020, PL/CK/1-3T/2020, and PL/Dk/13-16C/2020 were closely related to a Poland strain (gammaCoV/CK/Poland/G101/2016) (accession number MK576139) with over 95% identity and formed a common cluster within the Genotype I-23 (GI-23). PL/CK/17-20T/2020, PL/Ck/1-3T/2020 showed 99% homology between each other. While nucleotide sequence obtained from chicken PL/CK/1-5/2020 was closely related to a Nigerian isolate NGA/324/2006 with accession number FN182272 and clustered on the phylogenetic tree within the genotype 14 (GI-14) with 85% nucleotide similarity (Figure I). The obtained sequences of the *S1* gene of PL/CK/17-20T/2020, PL/CK/1-3T/2020, PL/CK/13-16CL/2020 and PL/CK/1-5CL/2020 were submitted to GenBank under accession number: OK072694, OK072695, OK072696, and OK072697.

**Figure I.**
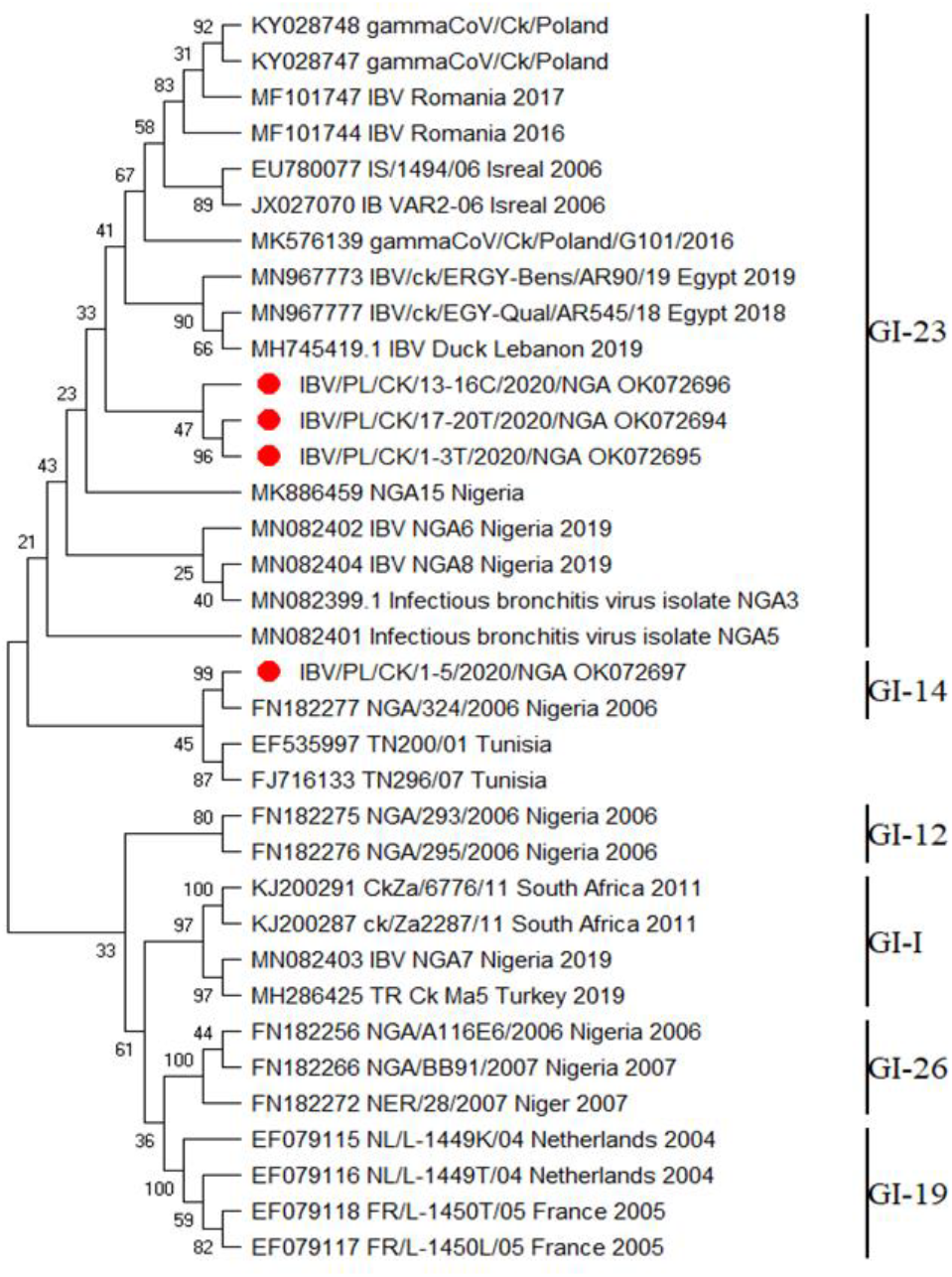
Phylogenetic tree of IBV based on the alignment of the nucleotide sequences of the *S1* gene. Sequences were aligned using MUSCLE with default settings in BioEdit (version 7.2.5.0). The phylogenetic tree was constructed by the Neighbor-Joining method with 1,000 bootstrap replicates. The evolutionary history was inferred using the Neighbor-Joining method. IBV detected in Jos LBM is labelled in red colour.

## Discussion

The present study reports for the first time the molecular detection and characterization of two different distinct genotypes of IBV (GI-14 and GI-23) LBMs in Jos. The LBMs plays an important role in the epidemiology of avian viral diseases (Bitrus *et al*., 2020; Sulaiman *et al*., 2021). This due to the known fact it is a major place enabling dissemination of major viral diseases. They also provide conditions for viral potential reassortment because of the high population densities and the mixture of poultry, poultry pets, and wild birds that are often present (Sulaiman *et al*., 2021). LBMs with limited biosecurity measures might be an important reservoir where new IBV variants can emerge. Overall prevalence of 42% recorded in this study is higher than the 13.19% reported by Mungadi et al. (2019) in Sokoto, 18% reported by (Ducatez *et al*., 2009). Detection of IBV in this study contrast the report by (Ducatez *et al*., 2009) who reported that IBV infection is not common in Northern Nigeria and in Niger republic backyard poultry. Alp Onen and Ozgur, (2017) also reported prevalence of 27.5% IBV in Turkey. Prevalence of 38% was also reported by (Bitrus *et al*., 2020) in three State of North-western Nigeria. Higher molecular detection of 58.8% and 63.4% was reported by Roussan *et* al. (2009) in Jordan and by Selim *et al*. (2013) in Egypt respectively. The difference in the prevalence in these studies might be due to the sensitivity of the different molecular techniques carried out as RT-qPCR molecular diagnostic assay targeting the highly conserved UTR gene have better sensitivity than conventional RT-PCR in detecting IBV nucleic acids (Acevedo *et al*., 2013). The disparity may also be attributed to the difference in time of the sampling, species of birds sampled, differences in sample size, types of management, nutritional status of birds and geographical location all of which can also played a greater role in the difference. Association of LBMs with IBV do not significantly differ (P > 0.05) in the study. The rate of positivity in all species of birds sampled in this study suggest a widespread transmission of the virus to other species of birds other than the chicken and its ability to replicate in other avian specie without showing clinical signs (Bitrus *et al*., 2020; Cavanagh *et al*., 2002). In this study, more positive results were obtained from the cloacal (27%) than from the tracheal swabs (15%), this contradicts the reports of Mungadi *et al*. (2019) who reported more positive results from the tracheal swabs (63.64%) than from the cloacal swabs. This also contradicts the reports of Bande et al. (2017) who stated that the upper respiratory tract is the primary replication site for IBV and initial infection starts at the epithelium of Harderian gland down to lower respiratory tract before reaching urogenital and gastrointestinal tracts The reports of this study is in harmony with the previous reports by Bhattacharjee and Jones (1997) and De Wit *et al*. (1998), that when birds are sampled in the chronic stage of an IBV infection, the value of examining gastrointestinal tract (caecal tonsils or cloaca swabs) is higher than that of examining the trachea. Jones, (2010) and El behery *et al*. (2016), also reported that infectious bronchitis virus replicates in the gut for longer periods than in the respiratory tract and the infection of gastro enteric tissues generally does not manifest itself clinically, but persists for long periods and results in faecal virus excretion where it is shed for several weeks in faeces after infection. This might suggest why numbers of positive samples were recorded in the cloacal than the tracheal samples. Sampling route in relation to IBV was not statistically significant (P > 0.05). Infectious bronchitis is not a well-studied disease in Nigeria possibly because it is not given keen attention as other respiratory pathogens and more especially because it is usually masked by other infections (Emikpe, 2010). Therefore, as reported by Callison *et al*. (2006), rapid differentiation of IBV infection from other respiratory tract diseases like the Avian influenza, Newcastle disease, Infectious laryngotracheitis, Avian Mycoplasmosis, Avian Metapneumovirus is important so that appropriate measures can be taken on time. The phylogenetic analysis of the virus from this study indicates that IBV isolates are diversified into two distinct genotypes: GI-14 and GI-23. Isolate IBV/PL/DK/1-3T/2020, IBV/PL/CK/1-3T/2020 and PL/CK/17-20T/2020 clustered together with the Polish strains (gammaCoV/CK/Poland/G101/2016) with 94-96% identity and other strain from the Middle East and North Africa in the GI-23 genotype. The GI-23 lineages are IBV variants circulating in the Middle East. This lineage continues to spread and poses a major challenge not only in the Middle East but also in many countries in Africa, Asia and Europe (Khataby *et al*., 2016). The GI-23 genotype was first isolated in West Africa in 2013 with the isolation of strains NGA3 (MN082399) and NGA8 (MN082404) in Nigeria *(Houta et al*., 2021). The positive case in the current study might have been introduced to the Nigerian poultry as a result of international trading in commercial exchanges of poultry and poultry related products, uncontrolled movement of animals across borders (Sulaiman *et al*., 2021) or through contaminated wild or migratory birds from the neighbouring countries as wild migratory birds have been found to play a greater role in the introduction and spread of viral diseases in different regions (De Wit & Cook, 2019). The detection of the IBV strain closely related with the European strain (Poland) agrees with previous report by (Houta *et al*., 2021) that some IBV lineages that are endemic in Europe have been isolated in African countries, and vice versa. The strains IBV/PL/CK/1-5CL/2020 that clustered in the GI-14 genotype with the Nigeria strain NGA/324/2006 with 85% identity was reported in 2006 which confirms the continued circulation of this strain in the country. This study established the widespread of the avian infectious bronchitis virus (IBV) in wide species of birds. The study also showed that the IBV isolates circulating in the study were diversified from two distinct genotypes: GI-14 and GI-23. Three isolates were closely related to a strain from Europe Poland, suggesting a possible introduction and formed a common cluster within the GI-23. One isolate was closely related to a Nigerian isolate NGA/324/2006 that was previously reported, indicating continuous circulation of the virus and formed a common cluster within the GI-14.

## Conclusion

This study established the widespread of the avian infectious bronchitis virus in different species of birds. The study also showed that the IBV circulating in the study were diversified into two distinct genotypes: GI-14 and GI-23. This study, therefore, recommends effective control Meas ures to mitigate the spread of the virus and continuous surveillance to identify the current circulating strain for possible local vaccine development to control the disease in the study area and the country at large was advocated. The results of this study must be seen in the context of several limitations that may be resolved in further studies. Initially, rather than screening each sample separately, they were combined into a pool. Second, there were not many samples collected from the different species in this study. A greater number of samples and bird species should be evaluated in future studies

## Acknowledgement

The authors acknowledge the scientific and technical support of the staff of the Regional Laboratory for Animal Influenza and Transboundary Animal Disease, National Veterinary Research Institute (NVRI), Vom. This project did not received any funding but was supported enormously with all reagents, primers, probes and all consumables needed in carrying out this research by NVRI, Vom, Nigeria.

## Conflict of Interest

The authors declare that there is no conflict of interest

## Notes

### Competing Interest Statement

The authors have declared no competing interest.

